# T-cell derived extracellular vesicles prime macrophages for improved STING based cancer immunotherapy

**DOI:** 10.1101/2023.01.09.523220

**Authors:** Aida S. Hansen, Lea S. Jensen, Kristoffer G. Ryttersgaard, Christian Krapp, Jesper Just, Kristine R. Gammelgaard, Kasper L. Jønsson, Mogens Johansen, Anders Etzerodt, Bent W. Deleuran, Martin R. Jakobsen

## Abstract

A key phenomenon in cancer is the establishment of a highly immunosuppressive tumor microenvironment (TME). Despite advances in immunotherapy, where the purpose is to induce tumor recognition and hence hereof tumor eradication, the majority of patients applicable for such treatment still fail to respond. It has been suggested that high immunological activity in the tumor is essential for achieving effective response to immunotherapy, which therefore have led to exploration of strategies that triggers inflammatory pathways. Here activation of the stimulator of interferon genes (STING) signaling pathway has been considered an attractive target, as it is a potent trigger of pro-inflammatory cytokines and type I and III interferons. However, immunotherapy combined with targeted STING agonists has not yielded sustained clinical remission in humans. This suggest a need for exploring novel adjuvants to improve the innate immunological efficacy. Here, we demonstrate that extracellular vesicles (EVs), derived from activated CD4^+^ T cells (T-EVs), sensitizes macrophages to elevate STING activation, mediated by IFNγ carried on the T-EVs. Our work support that T-EVs can disrupt the immune suppressive environment in the tumor by reprogramming macrophages to a pro-inflammatory phenotype, and priming them for a robust immune response towards STING activation.

## Introduction

The immune system, in particular activated T cells, are important for controlling tumor growth. However, tumors are often characterized with poor infiltration of activated T cells, or presence of dysfunctional T cells incapable of killing tumor cells [1]. Tumor associated macrophages (TAMs) are one of the most abundant immune cell populations in solid tumors, [2, 3] and are essential for regulating anti-tumoral T cell responses [4]. The phenotype and functions of TAMs are highly plastic. However, they can broadly be classified as either anti-tumorigenic or pro-tumorigenic. The majority of TAMs in solid tumors are pro-tumorigenic and play a role in inducing the immune suppressive tumor microenvironment (TME), favoring tumor progression, cancer cell invasion and metastasis [5]. Therapeutic reprogramming TAMs to an anti-tumorigenic phenotype is, therefore, a highly attractive strategy to improve current anti-cancer immunotherapy [6].

Activation of the stimulator of interferon genes (STING) signaling pathway is an effective trigger of type I Interferon and pro-inflammatory cytokines, supporting activation of the host immune system and polarization of macrophages to an anti-tumorigenic phenotype [7, 8]. Activation of the STING pathway involve multiple factors, but most importantly, the enzyme cGAS detect accumulated cytosolic DNA, which result in the production of cyclic-GMP-AMP (cGAMP). Binding of cGAMP to STING initiates a signaling cascade through TBK1, leading to downstream activation of the transcription factors IRF3 and NF-kB [9-12]. This support upregulation of type I interferon genes, and various cytokines and chemokines [9, 13, 14]. In addition to trigger macrophages, STING activation has also shown to support infiltration and activation of cytotoxic T cells in the TME [15-17] leading to increased tumor control in various tumor models [18-21]. However, immunotherapy targeting STING has not yielded sustained clinical remission in humans [22, 23], which may suggest a need for exploring novel adjuvants to improve the innate immunological efficacy.

Extracellular vesicles (EVs) have emerged as a novel mechanism of cellular communication and play a key role in regulating antitumor immune responses [24, 25]. EVs are small enveloped vesicles ranging in size from 30-1000 nm. They contain several biologically active molecules comprising proteins, lipids, and nucleic acids, to modulate the function of recipient cells [26-28]. They are produced by all cells and formed either by outward budding from the cell membrane as microvesicles or formed inside the cells in endosomes and released as exosomes [29-31].

Importantly, during the interaction of T cells with antigen-presenting cells (APCs), the T cells release high numbers of EVs [32] in a polarized manner at the interface with the APC [33]. These EVs are transferred unidirectionally to the APCs [34] to enhance their function [35, 36]. In addition, EVs from CD4^+^ T cells have also been demonstrated to promote antigen-specific B cell responses [37]. T-cell derived EVs contain, among other molecules, cytokines [38] and RNA fragments [39-41], and may therefore theoretically have the capacity to regulate surrounding immune cells by both direct receptor activation and by regulating gene expression. Some suggest T-cell derived EVs carry DNA capable of activating STING signaling in the APCs [35]. However, there is still very limited knowledge on how such T-cell derived EVs in fact modulates the immune response and in particular macrophages in the context of cancer [42].

In the present study, we demonstrate that EVs derived from activated CD4^+^ T cells modulate the function of macrophages and in particular prime the signaling capacity of the cGAS-STING pathway, supporting enhanced production of type I IFN and T-cell chemotaxic cytokines. We demonstrate that EVs from activated CD4^+^ T cells can be utilized as an adjuvant to improve the efficacy of *in vivo* STING agonist therapy for cancer.

## Results

### CD4^+^ T-cell derived EVs sensitize macrophages to STING activation

The ability of CD4^+^ T-cell derived EVs (T-EVs) to modulate activation of the innate immune system was examined by exploring different pattern recognition pathways in macrophages after priming with T-EVs. As some reports that T-EVs may carry surface-associated DNA [35], we consequently, and throughout the entire study, pre-treated T-EVs with DNase I prior to stimulation, to prohibit in-direct STING activation. THP-1 cells primed with T-EVs was exposed to stimulation with the STING ligand 2’,3’-cGAMP (cGAMP), the TLR3/RIG-I ligand poly(I:C) or the TLR4 ligand LPS, and assessed for cytokine production after 20 hrs. Priming with T-EVs resulted in a significantly enhanced production of type I IFN and IL-6 in response to activation by STING and TLR4. By contrast, priming did not affect the response to TLR3/RIG-I stimulation (**Fig. 1A, B**). Next, we showed that the priming effect of T-EVs in THP-1 cells was controlled in a cGAMP-concentration dependent manner. Interestingly, primed THP-1 cells were able to produce robust type I IFN and CXCL-10 at even very low cGAMP concentrations as compared to non-primed cells (**Fig. 1C, D**). This priming effect was not restricted to THP-1 cells only, as primary human monocyte-derived macrophages (MDMs) primed with T-EVs also gave significantly higher cytokine production to low-level dosage of cGAMP (**Fig. 1E, F**). Importantly, treating THP-1 or primary macrophages with T-EVs alone did not result in production of type I IFN, suggesting that T-EVs do not directly activate STING signaling (**Fig. 1E, F**). Finally, we evaluated the dose-dependent effect of T-EV particles needed to support priming of the STING pathway. Here, we found that at least 5 × 10^7^ particles were needed for priming of the cells (**Fig. 1G**). To summarize, our data indicate that EVs derived from activated CD4^+^ T cells are able to modulate the function of macrophages and in particular renders the STING pathway more sensitive to cGAMP stimulation without directly activating STING signaling.

**Figure 1.**
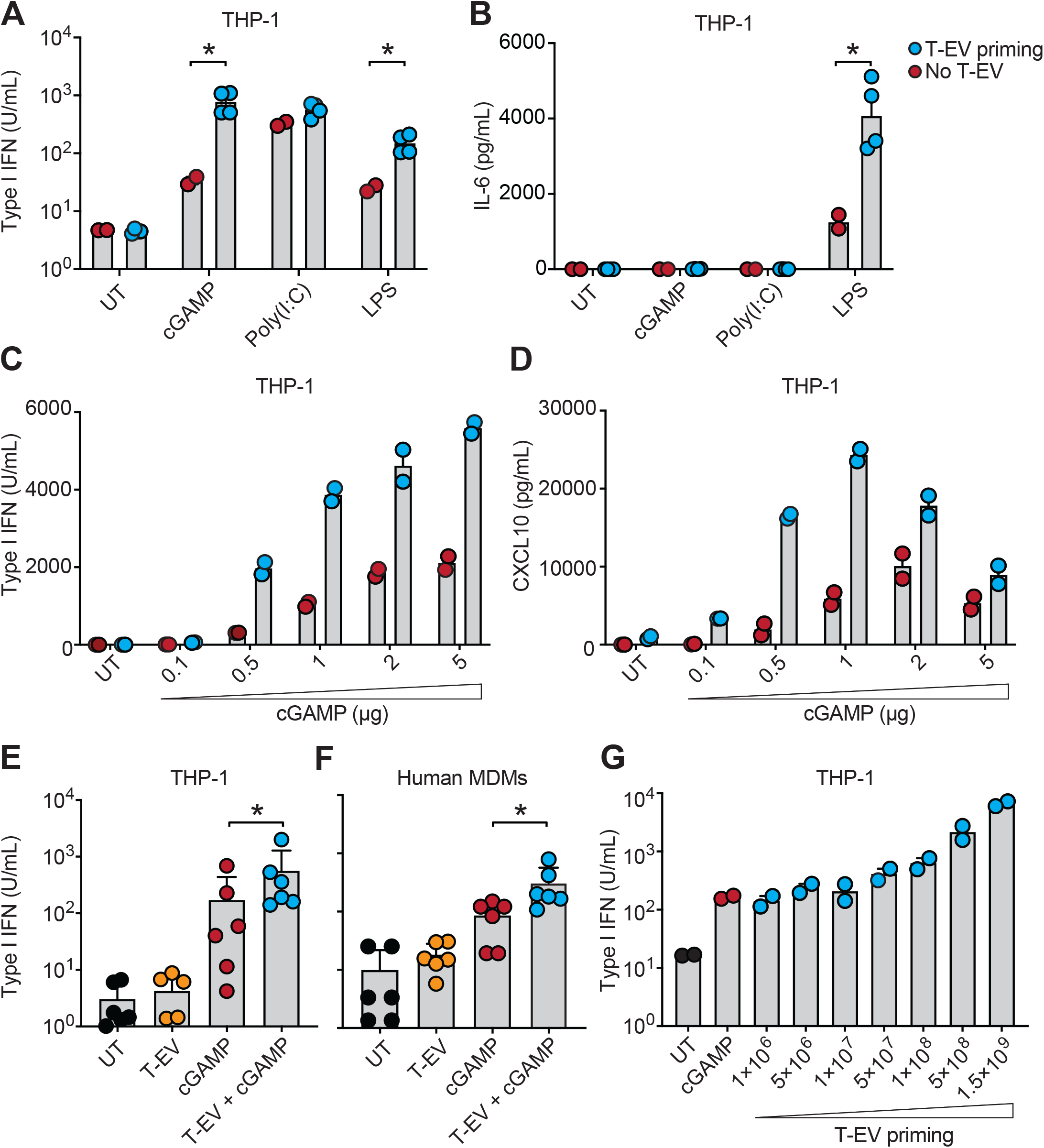
CD4^+^ T-cell derived EVs sensitizes macrophages to STING activation. **A)** and **B)** THP-1 cells were treated with T-EVs (1.5-2 × 10^9^) or EV-free media for 1 hr prior to stimulation with either cGAMP (0.5μg), Poly(I:C) (0.1μg) or LPS (0.5μg/mL). After 20 hrs of stimulation, the cell culture supernatant was analyzed for secretion of A) functional type I IFN and B) IL-6. Data show mean +SEM and individual replica of T-EVs from 2 distinct T cell donors, each in duplicates. *P < 0.05; unpaired t-test (two-tailed). **C)** and **D)** THP-1 cells were treated with T-EVs (3 × 10^9^) or EV-free media for 1 hr prior to stimulation with increasing amounts of cGAMP as indicated. After 20 hrs of stimulation, the cell culture supernatant was analyzed for secretion of C) functional type I IFN and D) CXCL-10. Data show mean +SEM and individual replica and are representative of 2 independent experiments. **E)** THP-1 cells and **F)** human monocyte derived macrophages (MDMs) were treated with T-EVs (3 × 10^9^) or EV-free media for 1 hr prior to stimulation with cGAMP (0.5μg). The production of type I IFN was determined after 20 hrs of stimulation. Data in E) shows mean +SD of 6 independent experiments and data in F) shows mean +SD of 6 different experiments all with T-EVs from distinct T cell donors. Each datapoint indicate mean value of duplicates. *P < 0.05; Wilcoxon test (two-tailed). **G)** THP-1 cells were treated with increasing amounts of T-EVs or EV-free media for 1 hr prior to stimulation with cGAMP (0.5μg) as indicated. The production of type I IFN was determined after 20 hrs of stimulation. Data show mean +SEM and individual replica and are representative of two distinct T-EV donors. UT = untreated, and received only EV-free media and lipofectamine. T-EV = EVs derived from activated CD4^+^ T cells.

### T-EVs modulate STING signaling independent of cGAS and intravesicular cGAMP

Importantly, DNase I treatment of T-EVs do not remove potential contaminating DNA inside the particles. Thus, to exclude that such mechanism could be a result of the priming effect, we used cGAS-deficient THP-1 cells as well as a STING-deficient control (**Fig. 2A, B**) generated by CRISPR-Cas9 editing [43]. The KO cell-lines responded similarly to PolyI:C stimulation (**Fig. 2C**) but upon priming with T-EVs and following cGAMP stimulation, we observed that cGAS-deficient THP-1 cells responded equally well as WT THP-1 cells (**Fig. 2D**). This support that the priming of macrophages by T-EVs is not mediated by intra-vesicular DNA. In parallel, priming of STING-deficient THP-1 cells with T-EVs and stimulating with cGAMP did not result in any IFN production (**Fig. 2D**). Potentially, T-EVs may also carry fractions of cGAMP, if such are produced by activated T cells. To rule out this scenario, we conducted a mass spectrometry-based detection assay of cGAMP in the T-EVs, but was unable to measure any detectable levels of cGAMP (**Fig. 2E**). Together, our data suggest that the function of T-EVs on sensitizing STING signaling is not mediated by neither DNA nor cGAMP inside the T-EVs.

**Figure 2.**
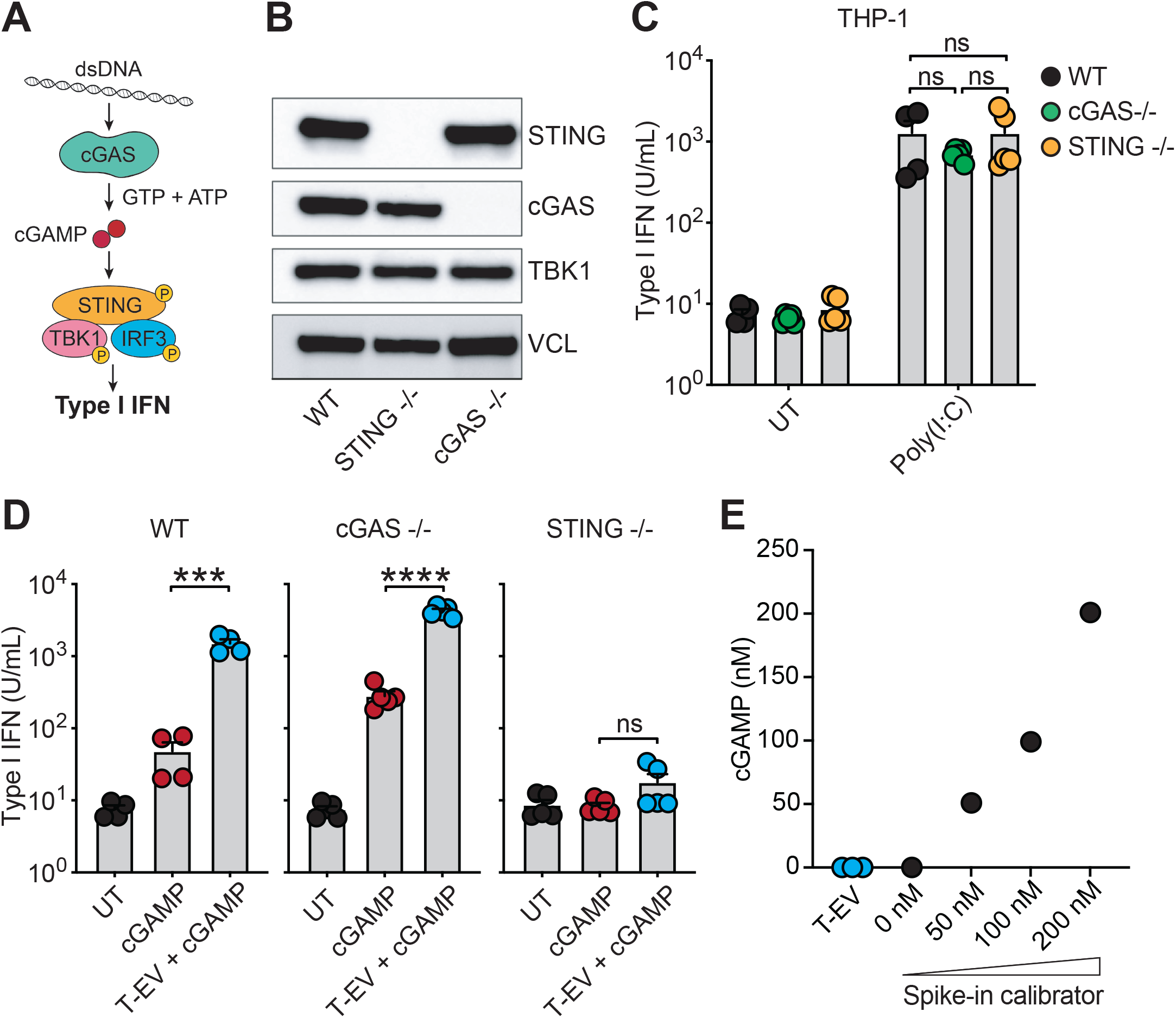
T-EVs modulate STING signalling independent of cGAS and intravesicular cGAMP. **A)** Schematics of the cGAS-STING pathway. **B)** Western blot analysis of cell lysates from WT, cGAS-/-, and STING-/- THP-1 cells. Vinculin (VCL) was used as a loading control. Data represents 1 experiment. **C)** WT, cGAS-/-, and STING-/- THP-1 cells were stimulated with Poly(I:C) (0.1μg). The production of type I IFN response was determined after 20 hrs of stimulation. Data show mean +SEM and individual replica of 2 independent experiments in duplicates or triplicates. ns = non-significant; unpaired t-test (two-tailed). **D)** WT, cGAS-/-, and STING-/- THP-1 cells were stimulated with T-EVs (3 × 10^9^) or EV-free media for 1 hr prior to stimulation with cGAMP (0.5μg). The type I IFN production was determined after 20 hrs of stimulation. Data show mean +SEM and individual replica of 2 independent experiments in duplicates or triplicates. ***P < 0.001; ****P < 0.0001; unpaired t-test (two-tailed). **E)** T-EVs were analyzed by mass spectrometry for presence of cGAMP. Data show cGAMP concentration in each of three distinct T-EV samples as well as in samples spiked-in with indicated amount of cGAMP. WT = Wild Type. UT samples in C) and D) are identical.

### T-EVs prime THP-1 cells for enhanced STING signaling

To further investigate the mechanism by which T-EVs prime the STING pathway, we examined potential changes in STING expression and function. After T-EV priming we did not observe any changes in *STING* mRNA expression compared to untreated cells (**Fig. 3A**). Next, we conducted a time-kinetic stimulation experiment with cGAMP. Cells primed with T-EVs resulted in a robust Type I IFN production as early as 4 hours after cGAMP was added to the culture (**Fig 3B**). We further examined if removing T-EVs from the THP-1 cell cultures prior to cGAMP stimulation would still support STING activation. Indeed, we observed that T-EV priming, independent of time prior to cGAMP stimulation, resulted in significant enhanced type I IFN production (**Fig. 3C**). This suggest that priming with T-EVs not only make cells more sensitive to low-level dosage of cGAMP, but also induce a more rapid initiation of the signaling pathway.

**Figure 3.**
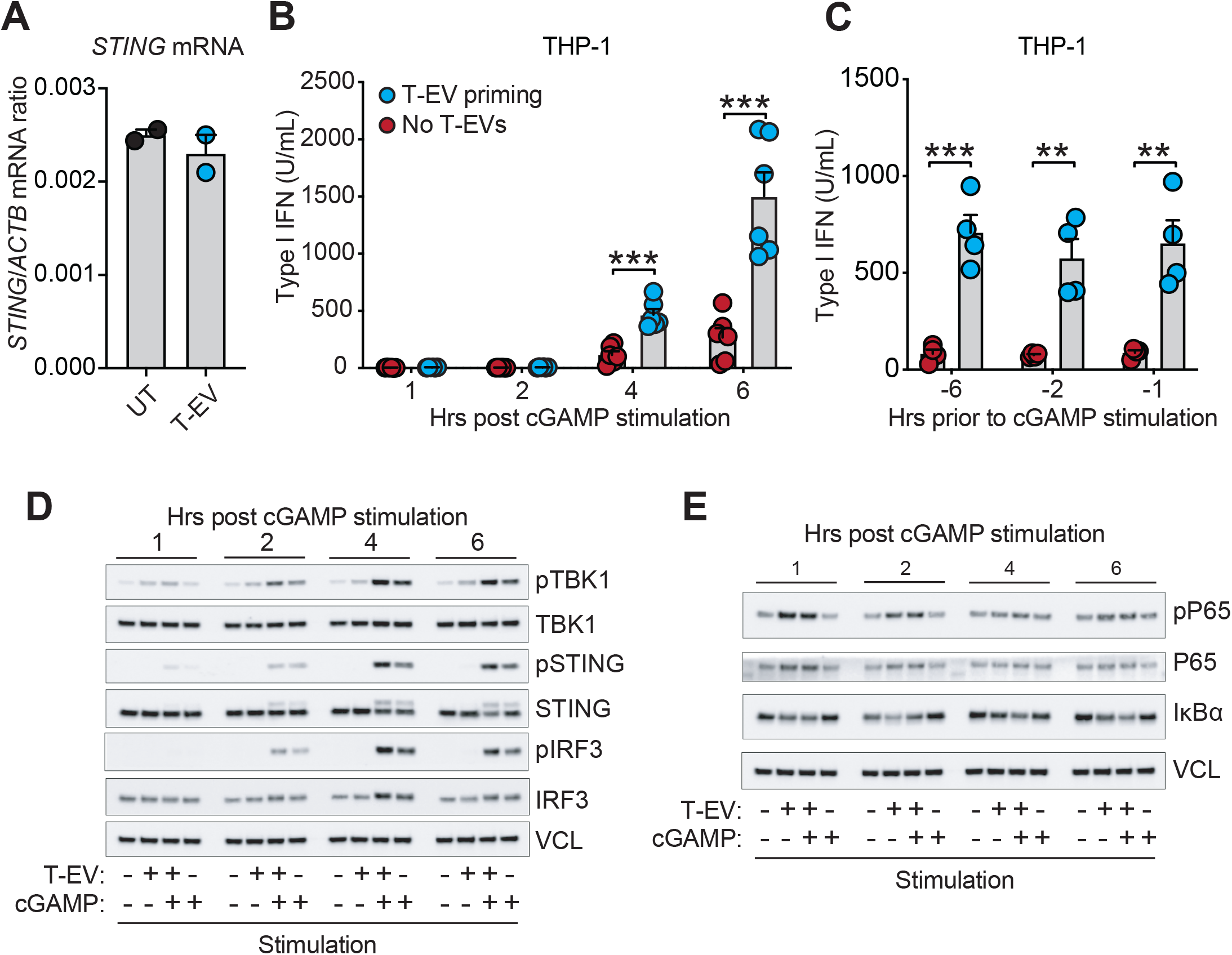
T-EVs prime THP-1 cells for enhanced STING activation. **A)** THP-1 cells were stimulated with T-EVs (3 × 10^9^) or EV-free media. After 6 hours of stimulation, the mRNA expression of STING was determined. Data shows mean +SEM and individual duplicates and is representative of 2 independent experiments. **B)** THP-1 cells were stimulated with T-EVs (3 × 10^9^) for 1 hr prior to stimulation with cGAMP (0.5μg). At indicated time-points after cGAMP stimulation the supernatant was harvested and the production of type I IFN response was determined. Data shows mean +SEM and individual replica of 3 independent experiments each in duplicates. ***P < 0.001; unpaired t-test (two-tailed). **C)** THP-1 cells were stimulated with T-EVs (1 × 10^9^) or EV-free media at the indicated time-points prior to cGAMP stimulation (0.5 μg). Right before cGAMP stimulation, the T-EVs were washed out. After 20 hrs of cGAMP stimulation, the supernatant was collected and the production of type I IFN was determined. Data show mean +SEM and individual replica of two independent experiments. ***P < 0.001; **P < 0.01; unpaired t-test (two-tailed). **D)** and **E)** THP-1 cells were stimulated with T-EVs (3 × 10^9^) or EV-free media for 1 hr prior to stimulation with cGAMP (0.5 μg). At indicated time-points after cGAMP stimulation, the cells were harvested. Expression and phosphorylation of indicated proteins were analyzed by Western blotting. Data are representative of 3 independent experiments. VCL was used as a loading control. VCL in D) and E) are identical.

To better understand how T-EV priming modulates response to STING activation, we investigated the early signaling pathway events upstream of IFN and cytokine gene expression, using western blotting analysis. T-EV priming alone did not lead to any clear signals in phosphorylation of STING and IRF3, though a minor increased intensity was observed for TBK1 (**Fig. 3D)**. However, comparing cGAMP stimulation of unprimed and primed cells, clearly showed more intense phosphorylation bands for T-EV primed cells (**Fig. 3D**). Furthermore, both pTBK1 and pSTING was increased already at 1 hour after cGAMP stimulation, supporting our earlier results suggesting a more rapid pathway activation. Activation of the STING pathway has also been reported to engage the IKK-NF-kB signaling pathway. Therefore, we next examined activation of p65, a component of the NF-kB complex, following T-EV and cGAMP stimulation. We observed both increased and more rapid phosphorylation of p65 upon priming with T-EVs prior to cGAMP stimulation compared to cGAMP stimulation alone (**Fig. 3E**). Also, the data indicated that T-EV priming alone, resulted in early phosphorylation of p65 which was sustained for up to 2 hours after priming (**Fig. 3E**). In all, these results suggest that the priming effect of T-EVs is a rapid event, and can be sustained in stimulated cells for long periods after the priming event happened.

### EVs from primarily activated CD4^+^ T cells enhances STING signaling

EVs are produced continuously from T cells, whether they are activated or not. Thus, to investigate whether STING sensitization was induced by T-EVs from CD4^+^ T cells in general, we isolated T-EVs from CD4^+^ T cells either **i)** stimulated with anti-CD3/anti-CD28 together with low-dosage of IL-2; or **ii)** left in the presence of only low-dosage IL-2. Activation status of the CD4^+^ T cells at the time of collection of T-EVs, was assessed by surface expression of CD69, CD134 and CD40L (**Supplementary fig. 1A-F**). We found that activation of CD4^+^ T cells was followed by an increased release of T-EVs into the cell culture supernatant (**Fig. 4A**). On the other hand, no difference in the size distribution between T-EVs from activated and non-activated CD4^+^ T cells was observed (**Fig. 4B, C**). The majority of T-EVs from both activated and non-activated CD4^+^ T cells were between 50-200nm, indicating that they consist of a mixture of both exosomes and microvesicles. Western blot analysis of lysate from T-EVs and their corresponding cells of origin showed that the EV-marker CD9 was exclusively expressed in the EV-fraction, whereas HSP70 and the marker for apoptotic bodies, calreticulin, was primarily expressed by the cellular fraction (**Fig. 4D**). This confirmed the identity of EVs and indicated low presence of apoptotic bodies in the T-EV preparations. Importantly, when priming THP-1 cells with these two batches of T-EVs from multiple T-cell donors, we observed that only T-EVs from activated CD4^+^ T cells showed a significant priming of STING (**Fig. 4E**). Consistent with this, we further showed that phosphorylation of p65 was primarily induced by T-EVs from activated CD4^+^ T cells (**Fig. 4F**). These data support that activation of CD4^+^ T cells leads to secretion of a distinct type of T-EVs with specific content capable of sensitizing the STING pathway in macrophages.

**Figure 4.**
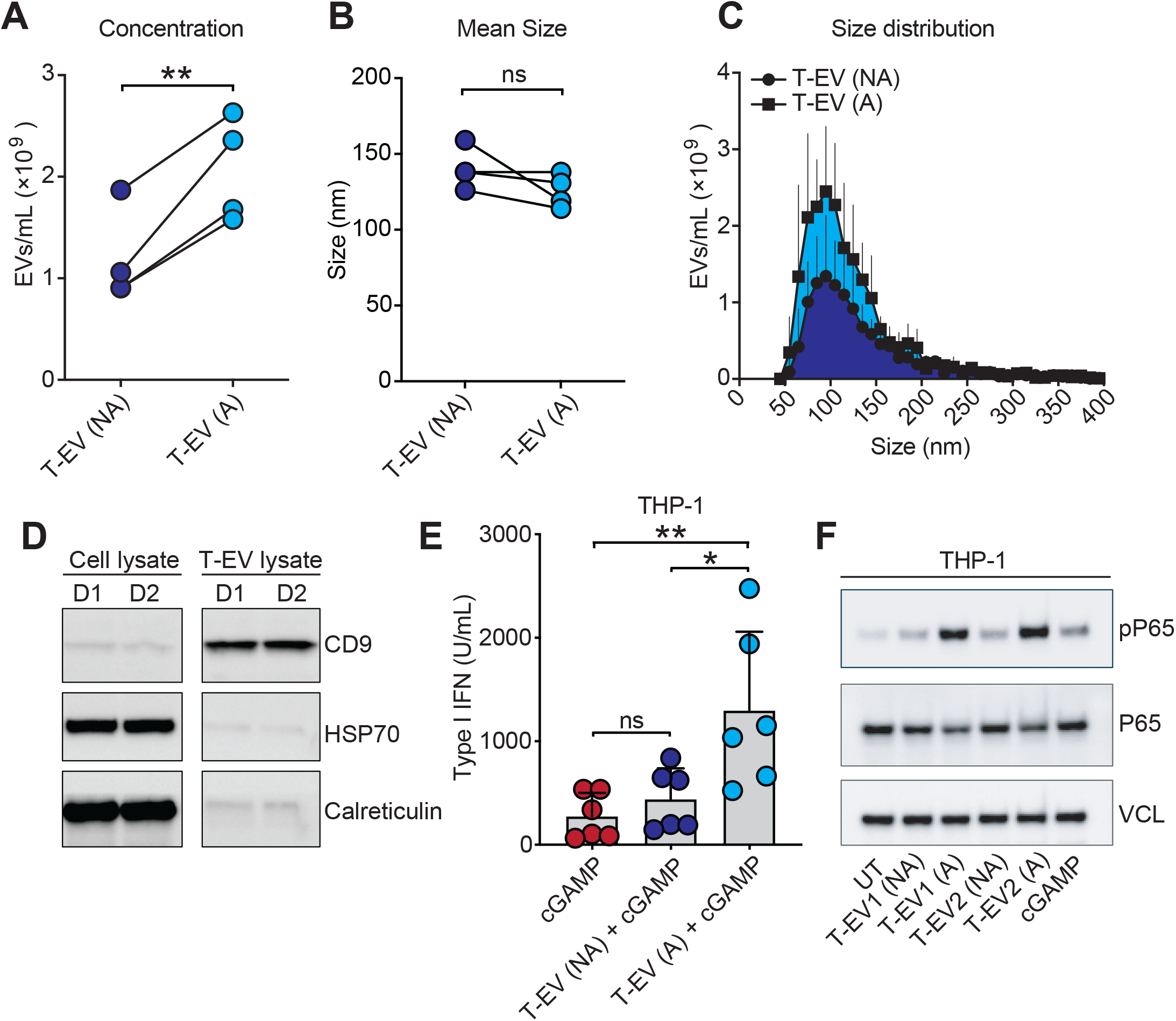
T-EVs from primarily activated CD4^+^ T cells enhances STING signaling. T-EVs were isolated from CD4^+^ T cells that were either activated (A) with anti-CD3 (1μg/mL) and anti-CD28 (1μg/mL), or left non-activated (NA) for 48hrs in presence of only IL-2 (10ng/mL). The T-EVs were analyzed by tunable resistive pulse sensing using a qNano for **A)** concentration and **B)** and **C)** size distribution. A) data shows T-EVs from each of 4 distinct donors. **P < 0.01; paired t-test (two-tailed). Data in B) show mean size of T-EVs from each of the 4 distinct donors. ns = not significant, paired t-test (two tailed). Data in C) show mean +SD of T-EVs from each of the 4 distinct donors. **D)** Western blot of paired cell lysate and T-EVs from two individual donors. **E)** THP-1 cells were treated with T-EVs (3 × 10^9^) from either A or NA CD4^+^ T cells for 1 hr prior to stimulation with cGAMP (0.5μg). The production of type I IFN was determined upon 20 hrs of stimulation. Data shows mean +SD and individual mean values of duplicates, from 6 different experiments. *P <0.05; **P < 0.01; paired t-test (two-tailed). **F)** THP-1 cells were stimulated with T-EVs (3 × 10^9^) from either A or NA CD4^+^ T cells for 1 hr prior to stimulation with cGAMP (0.5 μg). After 1 hr of stimulation, the cells were harvested for Western blot analysis. Data are from 1 experiment using T-EVs from two distinct donors. Vinculin (VCL) was used as a loading control. NA = non-activated. A = activated. ns = not significant.

### T-EVs transfer pro-inflammatory cytokines to enhance macrophage function

Various pro-inflammatory cytokines are known to trigger induction of NF-kB activation, and there is also evidence that cytokines can be transported between cells by EVs [38]. Prompted by our data, that T-EVs priming let to rapid pP65 activation, we therefore analyzed T-EVs for the presence of 10 inflammatory-associated cytokines using a highly sensitive multiplex immunoassay. Among these, we found that T-EVs from activated T cells contained both IFNγ, TNFα and IL-2 (**Fig. 5A**). Pre-treatment of THP-1 cells with these recombinant cytokines prior to cGAMP stimulation showed that mainly IFNγ and to a lesser extent TNFα enhanced the cGAMP-induced type I IFN production in a dose-dependent manner (**Supplementary fig. 2A, B**). Importantly, neither IFNγ, TNFα and IL-2 alone induced production of type I IFN. (**Supplementary fig. 2A-C**). Next, we compared the priming of THP-1 cells with a high-dose of recombinant IFNγ or TNFα compared to T-EV. These data showed that IFNγ alone was able to prime cells to a similar level as seen for T-EVs, whereas there was no further additive effect of combining IFNγ and TNFα (**Fig. 5B**). To test whether priming of the STING pathway was induced by cytokine content within T-EVs, we next blocked the receptors on THP-1 cells using antibodies recognizing IFNGR1 or TNFR1. To obtain efficient inhibition of IFNγ signaling we did also treat the T-EVs with anti-IFNγ before stimulating the cells with the T-EVs and subsequently cGAMP. Blocking TNFα signaling alone minorily affected type I IFN production (**Fig. 5C**) whereas blocking IFNγ signaling alone substantially abrogated the STING priming effect of T-EVs (**Fig. 5D)**. Blocking IFNγ and TNFα signaling simultaneously, did however, not further abrogate the STING priming effect of T-EVs (**Fig. 5E**). This suggest that T-EV-associated IFNγ may play an important role for priming STING signaling. These data further support that IFNγ may be carried on the surface of the T-EVs.

**Figure 5.**
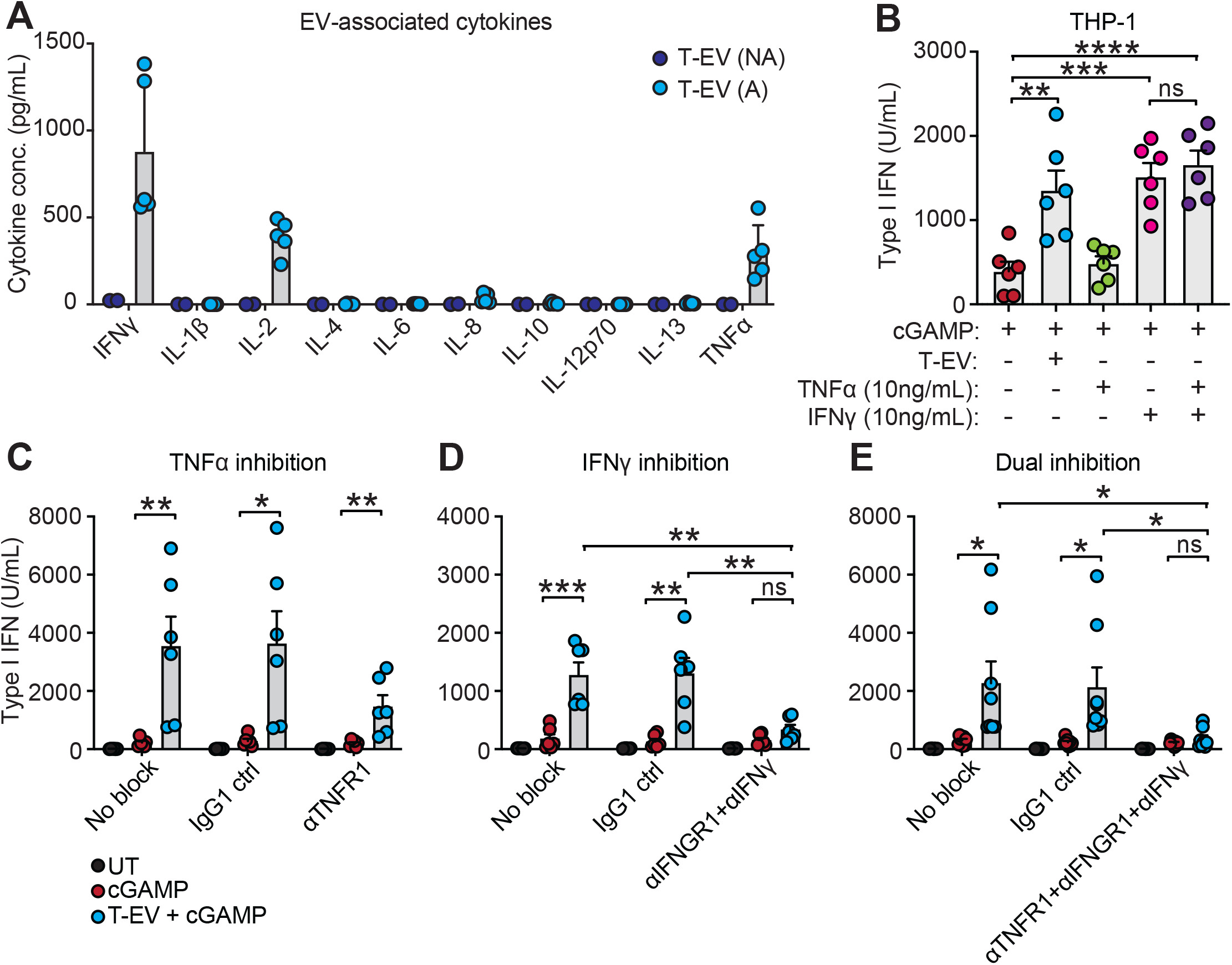
T-EVs transfer proinflammatory cytokines to enhance macrophage function. **A)** EVs from either activated (A) or non-activated (NA) CD4^+^ T cells were lysed in 2.5% Triton-X-100 and the presence of cytokines was measured using mesoscale or ELISA for IFNγ. Data show mean +SD and individual concentrations of n=5 (A) and n=2 (NA). **B)** THP-1 cells were stimulated with either T-EVs (1 × 10^9^), recombinant TNFα (10ng/mL) or recombinant IFNγ (10ng/mL) for 1 hr prior to stimulation with cGAMP (0.5 μg). The production of type I IFN was determined after 6 hrs of stimulation. Data show mean +SEM and individual replica from two independent experiments. **C)** THP-1 cells were treated with anti-TNFR1 (10μg/mL) or IgG1 (10μg/mL) for 30 min at 37°C. The cells were stimulated with T-EVs (1 × 10^9^) for 1 hr followed by stimulation with cGAMP (0.5μg). The production of type I IFN was determined after 6 hrs of stimulation. Data show mean +SEM and individual replica of three independent experiments. **D)** THP-1 cells were treated with anti-IFNGR1 (20μg/mL) or IgG1 (20μg/mL) for 30 min at 37°C. T-EVs were treated with anti-IFNγ (10μg/mL) or IgG1 (10μg/mL) and incubated for 15 min prior to use for stimulations. The cells were stimulated with the T-EVs (1 × 10^9^) for 1 hr followed by stimulation with cGAMP (0.5μg). The production of type I IFN was determined after 6 hrs of stimulation. Data show mean +SEM and individual replica of two independent experiments with T-EVs from three distinct donors in total. **E)** THP-1 cells were treated with a combination of anti-IFNGR1 and anti-TNFR1 as described for C) and D). Data show mean +SEM and individual replica of four independent experiments. *P < 0.05; **P < 0.01; unpaired t-test (two-tailed). Due to experimental setup, some of the datapoints for panel B) cGAMP and T-EV+cGAMP are identical to datapoints in panel D) and E) “No block”.

### T-EVs enhances the antitumoral function of cGAMP in vivo

So far, our results indicate that T-EVs can potentially function as an adjuvant for therapies targeting the STING pathway. To explore this further we moved to an *in vivo* syngeneic cancer mouse model. First, we confirmed cross-species activities, by exploring how T-EVs derived from activated murine CD4^+^ T cells functioned compared to our observations seen for human T-EVs. We primed murine bone-marrow derived macrophages (BMMs) with murine T-EVs prior to stimulation with a suboptimal dose of cGAMP. Similar to our observations in human macrophages, murine T-EVs enhanced the STING-induced production of IFN-beta (**Fig. 6A**). We further found that priming of BMMs with murine T-EVs alone induced phosphorylation of p65 (**Fig. 6B**). To confirm that our treatment was not toxic to cancer cells, we stimulated MC38 cells *in vitro* with murine T-EVs and cGAMP, but observed no effect on proliferation or cell death (**Supplementary fig. 3**).

**Figure 6.**
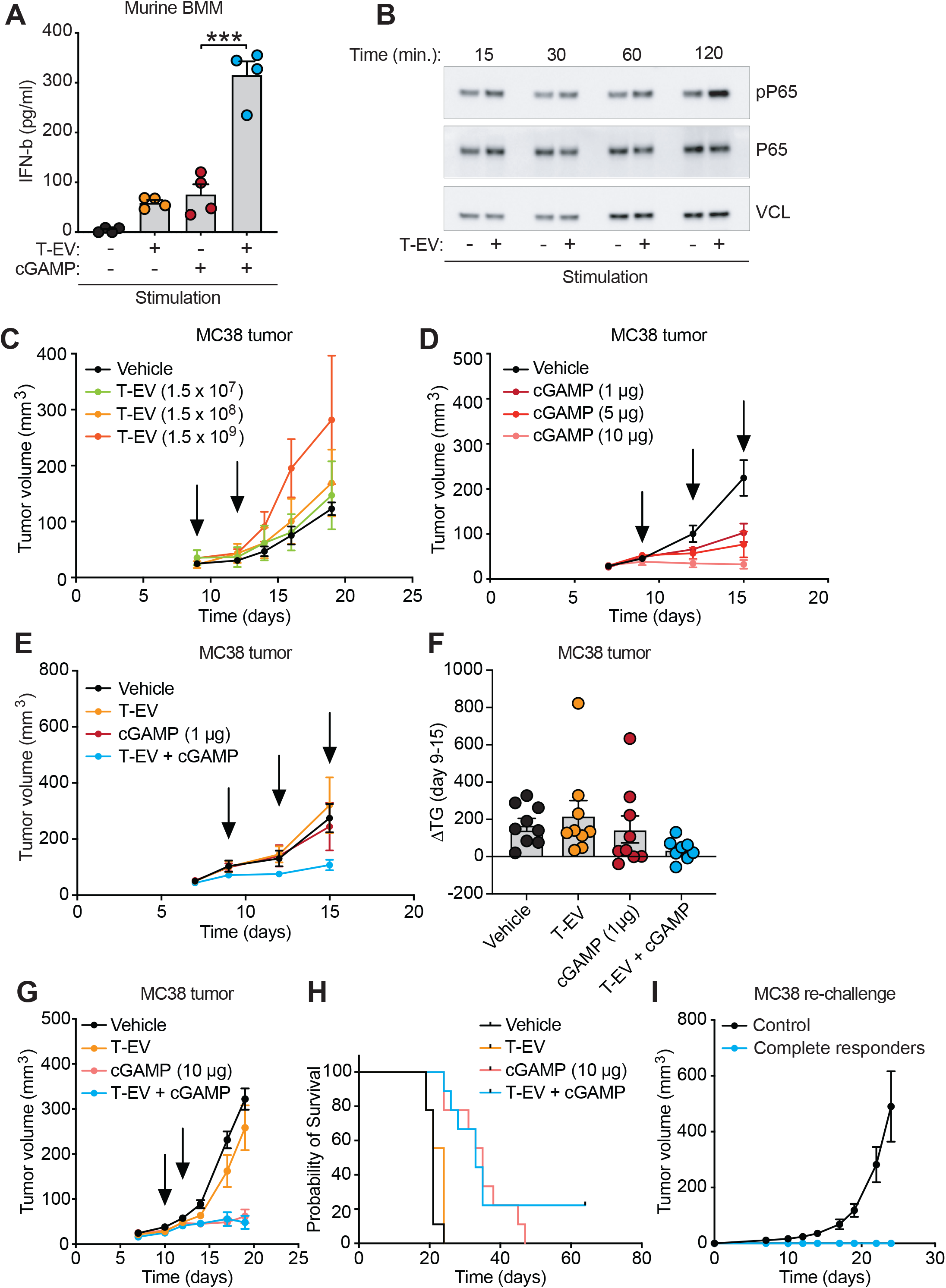
T-EVs enhance antitumor efficacy of cGAMP. **A)** Murine bone marrow derived macrophages (BMMs) were stimulated with murine T-EVs (1 × 10^9^) or EV-free media 1 hr prior to suboptimal cGAMP stimulation (0.05μg). The secretion of IFN-beta was determined after 20 hrs of stimulation. Data show mean +SEM and individual replica of two independent experiments each in duplicates. **B)** Murine BMMs were stimulated with murine T-EVs (1 × 10^9^) or EV-free media. At indicated time-points after T-EV stimulation, the cells were harvested and analyzed by western blotting for phosphorylation of P65. Data is from 1 experiment. VCL was used as a loading control. **C)** MC38 tumor bearing mice were treated with different amounts of murine T-EVs, administered intratumorally (IT) 2 times with 3 days interval as indicated with black arrows on the figurers. Treatment started on day 9 after tumor cell inoculation. Data show mean +/-SEM of tumor volume up to day 19 after tumor cell inoculation. n=4 in all groups. **D)** MC38 tumor bearing mice were treated with different amounts of cGAMP as indicated, administered IT 3 times with 3 days interval, as indicated with black arrows on the figurers. Treatment started on day 9 after tumor cell inoculation. Data show mean +/-SEM of tumor volume up to day 15 after tumor cell inoculation. n=6 in all groups. **E)** and **F)** MC38 tumor bearing mice were treated with either T-EVs alone (1.5 × 10^8^), cGAMP alone (1μg) or a combination. Mice were treated IT 3 times with 3 days interval as indicated with black arrows on the figurers, starting on day 9 after tumor cell inoculation. Data in E) show mean +/-SEM of tumor volume in mice treated with either Vehicle (n=9), T-EVs (n=9), cGAMP (n=9), or a combination of T-EVs and cGAMP (n=8) up to day 15 after tumor cell inoculation. Data in F) show difference in tumor growth from initiation of treatment on day 9 and until day 15, calculated as ΔTG. Bars indicate mean +/-SEM and individual value of ΔTG. TG = Tumor growth. **G)** and **H)** MC38 tumor bearing mice were treated with either T-EVs alone (1.5 × 10^8^), cGAMP alone (10μg) or a combination. Mice were treated IT 2 times with 2 days interval as indicated with black arrows on the figurers, starting on day 10 after tumor cell inoculation. Data in G) show mean +/-SEM of tumor volume in each group up to day 19 after tumor cell inoculation. Data in H) shows probability of survival up to day 64 after tumor cell inoculation. n=9 in each group. **I**) Mice that obtained complete tumor regression in G) was re-challenged with inoculation of MC38 cells subcutaneously in left flank. As controls was used C57B/6J mice (n=4). Data show mean +/-SEM of tumor volume in each group of either complete responders or control mice.

To investigate the antitumoral function of T-EVs, we next treated C57BL/6 mice bearing a subcutaneous MC38 colon adenocarcinoma with increasing amounts of murine T-EVs administered intratumorally in two repeated doses with 3-days interval. We found no anti-tumoral effect of the T-EVs alone (**Fig. 6C, Supplementary fig. 4**). Next, we titrated intratumoral administration of cGAMP to determine a suitable suboptimal dose, which we defined to be 1μg cGAMP leading to only a minor delay in tumor growth (**Fig. 6D, Supplementary fig. 5**). We then treated MC38 tumor-bearing mice with a combination of T-EVs and low-dosage of cGAMP in three repeated dosage with 3-days interval. After the first dosage, we observed a clear tendency to delayed tumor growth in the group recieving combination therapy compared to cGAMP monotherapy (**Fig. 6E, Supplementary fig. 6)**. When we calculated the tumor growth development during the treatment intervention, we saw a clear trend toward stagnated tumor growth in the combination group (**Fig. 6F**). To push for a better anti-tumoral effect, we next treated MC38-bearing mice with a combination of T-EVs and high-dose of cGAMP. Here we observed similar delay in tumor growth between mice recieving combined T-EV and high-dose cGAMP, and high-dose of cGAMP alone (**Fig. 6G, Supplementary fig. 7**). Intriguingly, in the group of mice recieving combinational treatment, we had 2/9 mice with complete tumor regression (**Fig. 6H**). Importantly, these mice were resistant to re-challenge with MC38 (**Fig. 6I**) indicating development of anti-tumor immunity. Altogether, our data indicate that treatment with T-EVs can prime the tumor microenvironment for a more potent immune response to cGAMP stimulation supporting improved control of tumor growth.

## Discussion

Activation of the STING pathway in the tumor microenvironment has shown profound anti-tumoral responses in preclinical models. However, it is unclear why these effects, to date, have not been achieved in clinical settings [23]. The usage of STING agonist as monotherapy and in combination with checkpoint inhibitors, lack to generate results that imply partly or complete responders. Potentially, this reflect that the dosage of STING agonists used, are either too low to induce signals or too high, which can ablate the essential immune cells in the TME needed for killing tumor cells [21]. Thus, there is a need to discover methods that expand the therapeutic index for proper STING activation in human settings.

One plausible explanation as to why STING agonists do not mount a powerful anti-tumoral response, is a physiolocal high threshold for activating the pathway in immune cells. A major question related to this is what endogenous level of cGAMP we can expect to measure in the TME. We currently lack compeling data demonstrating that endogenous levels of cGAMP accumulates in tumor tissue; and whether such concentrations are applicable for driving a potent STING-dependent immune response is difficult to answer. Some reports using murine tumor models suggest that blockage of cGAMP hydrolyzation in the extracellular space by the enzyme ENPP1 can support activation of STING [44]. Thus, ENPP1 inhibitors are considered promising therapeutics to enhance the level of extracellular cGAMP [45, 46]. Nonetheless, it is unresolved whether these levels of extracellular cGAMP overcomes the threshold of intracellular cGAMP needed for activating the STING pathway. In this study, we describe a promising mode-of-action for lowering the threshold for STING activation in immune cells. Cellular transmitters in form of extracellular vesicles (EVs) from activated CD4^+^ T cells were proved to prime macrophages and lower their responsiveness to even very small amounts of cGAMP. This mechanism of “alertness” resulted in both a rapid and strong cytokine induction profile, which sensitized tumors *in vivo* to mount effective tumor growth control at suboptimal dosage of STING agonists.

Conceptionally, the usage of adjuvants to support a strong and broad immunological response has been applied for decades in vaccine settings. Thus, applying such rationale for immunotherapy combination with STING agonists is an intriguing concept. There seem to be a delicate balance between robust activation of STING and induction of apoptosis, which needs to be considered in therapeutic settings. Work from Benoit-Lizon et al. and Larkin et al. suggest that the dosage of STING agonists to trigger type I IFN responses in various immune cells, will have detrimental effects on the viability of T cells [47, 48]. Thus, do we favor a powerful inflammatory cytokine response in the TME over the exsistance of potentially anti-tumoral lymphocytes? Our work propose that pre-activation of CD4^+^ T cells, which allow secretion of EVs that primes myeloid cells, can be beneficial prior to a direct STING activation. This could allow administration of STING agonists at dosages that both secure the presence of cytotoxic T cells but also elevates inflammatory cytokines in the TME.

EVs are continuosly secreted from cells and it is well appreciated that they can modulate multiple aspects of the TME, where their function closely reflect the characteristics of their cell of origin [25, 36, 49]. Thus, it is relevant to point out that in our study, we found that only EVs released from activated CD4^+^ T cells were able to sensitize STING signaling in macrophages. This raise the question whether EVs released from tumor infiltrating CD4^+^ T cells, which are often dysfunctional, may carry similar features or not? The usage of immune checkpoint inhibitors (ICI), that function by releasing the physiological break on T cell activation, could potentially be associated with elevated secretion of STING priming EVs. However, further work will be needed to address this.

Using a selective cytokine analysis, we found that EVs from activated T cells carried a series of known inflammatory signalling cytokines. Similar observations have been reported earlier, where a broad range of cell types seems to have the capacity to encapsulate cytokines within EVs [38, 50]. Interestingly, cytokines within EVs can both be surface bound and internalized, thereby mediating different signaling events to target cells. Our finding indicated that both IFNγ and TNFα was primarily surface bound, as blocking their designated receptors on macrophages prohibited the sensizatiation of the STING pathway. Multiple studies have described that tumors responding to ICI can be associated with active IFNγ signaling [51, 52], and the effect of IFNγ is associated with modulation of cancer cells and their activation of the STING pathway by increasing DNA damage and cGAMP production [53-55]. In parallel, IFNγ is also essential in activating and inducing a pro-inflammatory phenotype in macrophages [56-58]. However, the therapeutic application of IFNγ in cancer have been challenging because of the broad expression of the IFNGR and pleiotropic function of IFNγ in the TME [59]. EVs carrying cytokines on their surface may provide a targeted delivery of IFNγ to antigen-presenting cells in the TME and could influence the functional outcome and favor some of the key anti-tumorigenic functions of IFNγ. Thus, our work support a model where a combination of elevated cGAMP production within IFNγ-activated cancer cells with IFNγ- sensitized macrophages, are essential factors supporting modulation of an anti-tumoral STING-mediated immune response.

In conclusion, we show that IFNγ containing EVs derived from activated CD4^+^ T cells are able to sensitize macrophages, and potentially other immune cells, for enhanced STING signaling. We speculate that such EVs can disrupt the immune suppressive TME by reprogramming macrophages to a pro-inflammatory phenotype. In addition, they may render immune cells more sensitive to low-level STING agonist concentrations in the TME, supporting signaling pathway activation. Although clinical trials of STING agonists as cancer therapy showed mixed results, clinical development remains active [22]. We see that EVs from activated CD4^+^ T cells may be an exciting new avenue of therapy in combination with either ICI or other therapies supporting increased intratumoral cGAMP production, including radiotherapy or chemotherapy.

## Acknowledgement

This work was supported by Independent Research Fund Denmark (8026-00018B to A.S.H.); Novo Nordisk Foundation (Distinguished Innovator, NNF20OC0062825 to M.R.J.); Danish Cancer Society (R167-A10737-17-S2 and R246-A14695 to M.R.J., and R149-A10167-16-S47 to B.W.D.), Lundbeck foundation (R238-2016-2708 to M.R.J.), AUFF NOVA (AUFF-E-201 7-9-2 to B.W.D), and Gilead (Gilead Nordic Fellowship Programme 07591 to B.W.D). Support with NTA analysis was provided by Yan Yan, Interdisciplinary nanoscience center, Aarhus University, Denmark, and Kristian Juul-Madsen, Department of Biomedicine, Aarhus University, Denmark. Flow cytometry was performed at the FACS Core Facility, Aarhus University, Denmark. We kindly thank Ane Kjeldsen and Karin Skovgaard for lab assistance, and members of the Jakobsen and Deleuran labs for critical suggestions.

## Author contribution

Conceptualization, A.S.H., M.R.J., B.W.D; Methodology, A.S.H., M.R.J., A.E.; Investigation, A.S.H., L.S.J., K.G.R., C.K., J.J., K.R.G., K.L.J., A.E.; Writing – original draft, A.S.H.; writing – review and editing, A.S.H., M.R.J., B.W.D.; Visualization, A.S.H.; Funding acquisition, M.R.J., A.S.H., B.W.D.; Resources, M.R.J., B.W.D., M.J.; Supervision, M.R.J., B.W.D.; Project Administration, A.S.H.

## Materials and methods

### Isolation of human CD4^+^ T cells

Buffy coats from healthy donors were anonymously obtained from the blood bank at Aarhus University Hospital, Skejby, Denmark. Human peripheral blood mononuclear cells (PBMCs) were isolated from the buffy coats using Ficoll-Paque PLUS (GE Healthcare Bioscience). The PBMCs were cryopreserved in 40% heat-inactivated fetal bovine serum (FCS, Gibco), 50% RPMI-1640 (Sigma-Aldrich) and 10% DMSO (Sigma-Aldrich) and stored at −150 °C. CD4^+^ T cells were isolated from PBMCs by negative selection using EasySep Human CD4^+^ T Cell Isolation Kits (Stemcell) according to manufacturer’s instructions.

### Isolation of murine CD4^+^ T cells

The spleen was dissected from healthy C57BL/6 mice and a single cell suspension of splenocytes was prepared by pressing the spleens gently through a 70μm nylon cell strainer. CD4^+^ T cells were isolated from the splenocytes by negative selection using EasySep Mouse CD4^+^ T Cell Isolation Kits (Stemcell) according to manufacturer’s instructions.

### Generation of extracellular vesicles (EVs)

For generation of EVs the cells were cultured in medium containing EV-free FCS prepared by ultracentrifugation at 100,000 × g for 20 hours to remove all EVs from the serum. CD4^+^ T cells were activated *in vitro* resulting in release of high numbers of extracellular vesicles (EVs) into the cell culture supernatant. For activation of human CD4^+^ T cells, approximately 4-6 × 10^6^ isolated human CD4^+^ T cells/well were stimulated in a 6-well plate pre-coated with 1μg/ml αCD3 (clone UCHT-1, R & D System) and cultured in RPMI-1640 supplemented with 10% EV-free FCS, 10 mM HEPES (Gibco), 2mM glutaMAX (Gibco), 200 IU/ml Penicillin, 100 μg/ml Streptomycin, 1 μg/ml soluble αCD28 (clone CD28.2, BD Pharmingen) and 10 ng/ml recombinant human IL-2 (Roche). For activation of murine CD4^+^ T cells, approximately 4-6 × 10^6^ isolated murine CD4^+^ T cells/well were stimulated in a 6-well plate pre-coated with 1μg/ml αCD3 (clone 145-2C11, BD Pharmingen) and 1μg/ml and αCD28 (clone 37.51, BD Pharmingen) and cultured in RPMI-1640 supplemented with 10% EV-free FCS, 10 mM HEPES (Gibco), 2mM glutaMAX (Gibco), 1X non-essential amino acids (Gibco), 200 IU/ml Penicillin, 100 μg/ml Streptomycin, 50μM beta-mercaptoethanol and 10 ng/ml recombinant human IL-2 (Roche). Both human and murine CD4^+^ T cells were cultured for 46 hrs under humidified conditions at 37°C, 5% CO_2_.

### Flow cytometry analysis

CD4^+^ T cells were collected for flow cytometry analysis before and after activation as described above. 2 mio cells were stained using the following antibodies: anti-CD3-BV421 (BD Horizon), anti-CD4-FITC (BD Pharmingen), anti-CD45RA-V500 (BD Horizon), anti-CD69-BV786 (BD Horizon), anti-CD134-PE-Cy7 (BD Bioscience), and anti-CD40L-BB700 (BD Optibuild). LIVE/DEAD Fixable near-IR (Thermo Fisher Scientific) was used for live/dead discrimination. Samples was analyzed using the NovoCyte Quanteon and data processed in FlowJo.

### Isolation of EVs

EVs were isolated from the cell culture supernatant by differential centrifugation. In brief, the cell culture supernatant was centrifuged for 10 minutes at 1000 × *g* followed by 20 minutes at 2000 × *g* and 35 minutes at 10,000 × *g*. The EVs were then pelleted from the supernatant by centrifugation for 2 hrs at 100,000 × *g*. All centrifugation steps were performed at 4°C and the ultracentrifugation was done using a TST 41.14 rotor (Sorval). The EV pellet was resuspended in either PBS (Sigma-Aldrich), T cell media or physiological water (Sigma-Aldrich). To remove surface associated DNA the EVs were incubated with 10 units DNase I (Thermo Scientific) for 1 hr at room temperature.

### Nanoparticle tracking analysis

Throughout the entire study, the concentration of T-EVs was analyzed by nanoparticle tracking analysis (NTA) to adjust the amount of T-EVs used for functional studies. NTA was performed in scatter detection mode, using a NanoSight NS300 (Malvern Instruments) with a 405 nm laser. The EVs were diluted 100 times in PBS and measurements were performed in five times of 60s video captures of each sample with camera level 13 and detection thresshold 4-5 for all analysis. Data were analysed using the NTA software version 3.4.

### EV characterization

A qNano Gold (Izon Bioscience) equipped with a NP100 Nanopore (analysis range 50-330 nm, Izon Bioscience) for tunable resistive pulse sensing (TRPS) was used to determine size and concentration of the EVs. The EV samples were diluted in PBS and the EV samples were analyzed under identical settings - the same diluent, stretch (47mm), pressure (10Pa) and voltage (0.6mV) as used for CPC100 calibration particles (Izon Bioscience). The EV concentration and size distribution were determined using the “Izon control suite” software (Izon Bioscience).

### Mass Spectrometry for cGAMP analysis

Mass spectrometry analysis for presence of cGAMP within the T-EVs was done according to previously described [9]. In brief, 6-9.5 × 10^9^ EVs from activated human CD4^+^ T cells were resuspended in 1ml v/v 80%, 2% acetic acid. The samples were incubated for 15min at RT and added to NH_2_ SPE-columns pre-conditioned with first methanol and water. Columns containing the EV samples were washed with methanol and subsequently water and cGAMP was eluted with alkaline methanol (v/v 80% methanol, 5% NH_4_OH). The eluted samples were dried using a vacuum centrifuge and dissolved in 50μl 0.1% formic acid. 10μl of the samples were injected on a HSS T3 (2.1 × 100mm) LC-column. cGAMP was semi-quantified using LC-MS/MS with the transition *m/z* 338 > 152 in a positive ionization mode (ESI(+)). As external calibrators, corresponding blank samples were spiked with 0, 50, 100 and 200 nM cGAMP respectively and worked up as the remaining samples prior to analysis.

### Cell culture

All cells were cultured under humidified conditions at 37°C and 5% CO_2._ Human acute monocytic leukemia cell line (THP-1) was cultured in RPMI-1640 (Sigma-Aldrich) supplemented with 10% heat-inactivated fetal calf serum (FCS, Gibco), 200 IU/ml Penicillin, 100 μg/ml Streptomycin and 600 μg/ml glutamine (hereafter termed RPMI complete). Infection with mycoplasma was tested and ruled out on a regular basis by Eurofins. For all experiments, THP-1 cells were differentiated into phenotypically adherent macrophages by stimulation with 100 nM Phorbol 12-myristate 13-acetate (PMA, Sigma-Aldrich) in RPMI complete, for 24 hours. The medium was then refreshed with normal RPMI complete, allowing the cells to further differentiate an additional day. They were hereafter defined as macrophages. THP-1 clones harboring knock-out mutations in genes encoding STING and cGAS was previously described [43]. These cells were cultured and activated similar to WT THP-1 cells.

Human monocyte-derived macrophages (MDMs) were differentiated from PBMCs by culturing the cells in RPMI complete medium supplemented with 10% AB-positive human serum and 15 ng/ml M-CSF (PeproTech, 300-25-100UG). After 2 days of culturing, the medium was changed to Dulbecco’s Modified Eagle Medium (DMEM, Sigma-Aldrich) supplemented with 200 IU/ml Penicillin, 100 μg/ml Streptomycin, 600 μg/ml glutamine, 10% AB-positive human serum and 15 ng/ml M-CSF and the cells were allowed to further differentiate for additional 6-8 days.

Murine bone marrow was extracted from femur and tibia from 8-12 weeks old healthy C57B6/J mice by centrifugation, and cryopreserved at −150°C in 40% heat-inactivated fetal bovine serum (FCS, Gibco), 50% RPMI-1640 (Sigma-Aldrich) and 10% DMSO (Sigma-Aldrich). Bone marrow derived macrophages (BMMs) were differentiated by culturing bone marrow cells in presence of L929 SN containing the necessary M-CSF for differentiation. L929 SN was prepared by collecting the supernatant (SN) from confluent L929 cells. For differentiation, bone marrow cells were cultured in RPMI-1640 complete medium supplemented with 20% L929 SN. After 2 days the cells were refreshed with RPMI complete medium supplemented with 40% L929 SN and on day 4 the medium was changed to RPMI complete supplemented with 20% L929 SN. After 6 days the BMMs were fully differentiated and harvested using Accutase and seeded for further stimulation.

### Stimulation of cells in vitro

For stimulation of the cells *in vitro* were used 5 × 10^4^ cells/well in a 96-well plate of either PMA-differentiated THP-1 cells, MDMs or BMMs. The cells were seeded 1 day prior to stimulation allowing the cells to adhere to the bottom of the culture well. Upon stimulation, the culture medium was removed from the cells and 50 μl EV-free medium containing 1-6 × 10^9^ EVs, or 50 μl EV-free medium alone was added to the cells and they were incubated at 37°C for 1 hr. 2’3’-cGAMP (InvivoGen) (0.5 μg/well) or poly:IC (0.1μg/well) were mixed with Lipofectamine-2000 (Invitrogen) in a ratio of 1:1 according to manufacturer’s instructions. This mix was diluted with EV-free media and added to the cells in a volume of 100μl. The cells were incubated at 37°C for 20-24 hours if not otherwise indicated in the figures. Lipopolysaccharides (LPS) (Sigma-Aldrich) were diluted in EV-free media to a concentration of 0.5μg/mL and added to the cells in a volume of 100μl. Recombinant human (rh) Interleukin-2 (IL-2, Sigma-Aldrich), rhTNFα (Peprotech) and rhIFNγ (Peprotech) were diluted in EV-free media as indicated on the figurers, and added to the cells in a total volume of 50μl. The cells were incubated at 37°C for 1 hr and stimulated with 2’3’- cGAMP (0.5μg/well) as described previously. For inhibiting NF-kB signaling, cells were treated with 10μM ML120B (Tocris) for 30 min. at 37°C. For blocking cytokine signaling, cells were incubated with 10μg/ml human anti-TNFR1 (clone 16805) (R&D System), 20μg/ml human anti-IFNGR1 (clone GIR208) (R&D System) or human IgG1 (clone MOPC-21) (Biolegend) for 30 min. at 37°C. For blocking IFNγ signaling, the T-EVs was incubated with human anti-IFNγ (clone MD-1) (Biolegend) for 15 min. at 37°C.

### Real-time PCR

RNA was extracted from the cells using the RNeasy mini kit (Qiagen) according to manufacturer’s instructions. The concentration of the isolated RNA was determined using Nanodrop and 250ng RNA was converted into cDNA using the iScript cDNA synthesis kit (Bio-Rad) according to the manufacturer’s instruction. The cDNA was diluted by a final factor 25 for the real-time PCR analysis. Real-time PCR analysis of STING and ACTB was performed using the TaqMan Fast advanced Mastermix (Thermo Fisher Scientific) and the following TaqMan gene expression assays: Hs00736955_g1 (STING/TMEM173) and Hs01060665_g1 (ACTB). The analysis was done using the AriaMX Real-time PCR System (Agilent Technology).

### Functional type I IFN

Bioactive functional type I IFN was quantified in supernatants using the reporter cell line HEK-Blue IFN-α/β (Invivogen) according to manufacturer’s instructions. This cell line expresses secreted alkaline phosphatase (SEAP) under control of the IFN-α/β inducible ISG54 promoter. The cell line was maintained in DMEM supplemented with GlutaMax-I (Gibco, Life Technologies), 10% heat-inactivated FCS, 100 μg/mL streptomycin and 200 U/mL penicillin, 100 μg/mL normocin (InvivoGen), 30 μg/mL blasticidin (InvivoGen) and 100 μg/mL zeocin (InvivoGen). For measurement of functional type IFN, cells were seeded at 3 × 10^4^ cells/well in 96-well plates in 150 μL medium devoid of Blasticidin and Zeocin. The following day 50 μl of the supernatant for analysis were added to the HEK-Blue cells. SEAP activity was assessed by measuring optical density (OD) at 620 nm on a microplate reader (ELx808, BioTEK). The concentration was determined from a standard curve made with IFN-α (IFNa2 PBL Assay Science) ranging from 2 to 500 U/ml.

### Enzyme-linked Immunosorbent assay (ELISA)

Protein levels of the cytokines CXCL-10 and IL-6 in the cell culture supernatants were determined using the following ELISA kits: CXCL-10 and IL-6 (R&D System) according to manufacturer’s instructions. IFN-beta in the cell culture supernatants was determined using the mouse IFN-beta DuoSet ELISA kit (R&D System) according to manufacturer’s instructions. IFNγ in the T-EVs were determined using the Human IFNγ DuoSet ELISA kit (R&D System) according to manufacturers instruction. Prior to analysis, the T-EVs were lyzed in v/v 2.5% Triton-X-100, 0.05% Tween-20.

### Mesoscale V-Plex

1 × 10^9^ T-EVs from either activated (A) or non-activated (NA) human CD4^+^ T cells were lyzed using v/v 2.5% Triton-X-100, 0.05% Tween-20. This lysate was analyzed for presence of IFNγ, IL-1β, IL-2, IL-4, IL-6, IL-8, IL-10, IL-12p70, IL-13 and TNFα using V-plex (Meso Scale Discovery®, V-PLEX® Proinflammatory Panel 1 (human)) according to manufacturer’s instructions. Samples were diluted 1:2.

### Western blotting

Protein expression was analyzed by Western blotting. Cell lysis buffer was made by mixing Pierce RIPA buffer (Thermo Scientific) supplemented with 10 mM NaF, 2X complete protease cocktail inhibitor (Roche), 2X protease and phosphatase inhibitor (Pierce) in a 1:1 dilution with Laemmli Lysis Buffer (Sigma-Aldrich). Benzonase (Sigma-Aldrich) were added to the lysis buffer in a volume of 1μl/ml lysis buffer. For lysis of cells, 60μl of this lysis buffer was added to a single well in a 96-well plate. For lysis of T-EVs, 75μl of lysis buffer was added to the 100K T-EV pellet.

Protein concentration in the lysates was determined using the Pierce BCA Protein Assay Kit (Thermo Fisher Scientific) according to manufacturer’s instructions. The lysate was denatured at 95 °C for 5 minutes and separated on a Criterion Precast Gel, 4-12% Tris-HCl (Bio-Rad) with XT MOPS running buffer (Bio-Rad). The separated proteins were transferred to a Trans-Blot Turbo 0.2 μm PVDF membrane using the Trans-Blot Turbo Transfer System (Bio-Rad). Membranes were washed in TBS supplemented with 0.05% Tween-20 (TBS-T) and blocked in 5% fresh made skim-milk (Sigma-Aldrich) in TBS-T. Following antibodies were diluted in 1-5% BSA (Roche): anti-cGAS (Cell Signaling, D1D3G), anti-STING (Cell Signaling, D2P2F), anti-vinculin (Sigma Aldrich, v9131), anti-phospho-STING (Cell Signaling, D7C3S), anti-phospho-TBK1 (Cell Signaling, D52C2), anti-TBK1 (Cell Signaling), anti-phospho-IRF3 (Cell Signaling, D601M), anti-IRF3 (Cell Signaling, D83B9), anti-phospho-P65 (Cell signaling, 93H1), anti-P65 (Cell Signaling, D14E12), anti-IkBa (Cell Signaling, L35A5), anti-CD9 (Santa Cruz, C4), anti-HSP70 (Cell Signaling), and anti-Calreticulin (Cell Signaling, D3E6). The membranes were incubated with primary antibodies overnight at 4°C. The following secondary antibodies were diluted in 1% skim-milk: peroxidase-conjugated F(ab’)2 donkey anti-mouse IgG (H+L), peroxidase conjugated affinipure F(ab’)2 donkey anti-rabbit IgG (H+L), and peroxidase conjugated F(ab’)2 donkey anti-goat IgG (H+L) (all purchased from Jackson Immuno Research). The membranes were incubated with secondary antibodies for 1 hour at room temperature. The membranes were exposed using Clarity Western ECL Blotting Substrate.

### Cell proliferation assay

Proliferation of MC38 cells was measured using the CellTiter-glo 2.0 Assay (Promega) according to manufacturer’s instructions. In brief, 5000 MC38 cells/well were seeded in a 96-flat well plate the day before stimulation. The cells were treated in triplicates with T-EVs and cGAMP as previously described. As a control, the cells were treated with 0.1μM Doxorubicin hydrochloride (Sigma-Aldrich). Luminescent signal was determined after 48 hrs of stimulation.

### Mice

C57BL/6 mice from Janvier were housed in the animal facility at Department of Biomedicine, Aarhus University, under conditions according to the recommendations of The Animal Experiments Inspectorate under the Ministry of Environment and Food of Denmark. The Study was conducted in accordance with The Animal Ethics Council, license number: 2017-15-0201-01253. Female mice with an age of 7-9 weeks at initiation of the experiments, were used for all experiments.

### Generation of tumor xenografts

MC38 (Kerafast) colon adenocarcinoma cells were cultured in DMEM (Sigma Aldrich) supplemented with 10% FCS, 2mM Glutamine, 1000 U/ml penicillin and 1000 μg/ml streptomycin, 1X non-essential amino acids (Gibco). The cells were cultured under humidified conditions at 37°C and 5% CO_2_. The cells were split before they reached 100% confluency and were detached from the cell culture flask using 0.25 % trypsin (Gibco) in PBS with 0.02% EDTA (Invitrogen). To establish a tumor in the mice, 1 × 10^6^ MC38 cells in 50μl were injected subcutaneously into the right flank of the mice under isoflurane anaesthesia.

### In vivo treatment

The *in vivo* experiment was conducted at day 9-10 after tumor cell inoculation, when the tumors reached a size of 20-40 mm^3^. The mice were grouped randomly and treated intratumorally (IT), with 30μl containing either different amounts of 2’3’-cGAMP Vaccigrade (Invivogen), murine T-EVs diluted in physiological water (Sigma-Aldrich) or a combination. The tumor size was measured regularly using a caliper and the tumor volume was calculated using the formula: Tumor Volume (mm^3^) = 0.5233*L*W*H where, L = Length (mm); W = Width (mm); H = Height (mm). At the end of the experiments or when the tumor reached a size of 1000 mm^3^, the mice were euthanized.

### Statistics

Data are in general shown as bars indicating mean +SD or +SEM, and with points indicating each replica. Statistical analysis was performed with Graphpad Prism software. Comparison between different groups were analyzed by either paired Students t-test, unpaired Students t-test or Wilcoxon test, as indicated in the figure legends. All p-values are two-tailed. Significance was defined as p<0.05.

