## Supplementary material for "T-cell derived extracellular vesicles prime macrophages for improved STING based cancer immunotherapy": Suplemental information

#### Supplementary information

**Supplementary figure 1. CD4$^+$ T cells express activation-associated markers at time of T-EV harvest.** Human CD4$^+$ T cells were either activated (A) with plate-bound anti-CD3 and anti-CD28 or left non-activated (NA) for 48hrs in presence of only IL-2. The expression of surface markers associated with T-cell activation was assessed by flow cytometry. **A)** Representative density charts and **B)** frequency of CD69$^+$ cells, shown as percentage of live CD4$^+$ T cells. Data in B) show mean +SD of three distinct donors. **C)** Representative density charts and **D)** frequency of CD134$^+$ cells, shown as percentage of live CD4$^+$ T cells. Data in D) show mean +SD of three distinct donors. **E)** Representative density charts and **F)** frequency of CD40L$^+$ cells, shown as percentage of live CD4$^+$ T cells. Data in F) show mean +SD of three distinct donors.

**Supplementary figure 2. Recombinant IFNγ and TNFα prime STING signaling in THP-1 cells dose-dependent.** THP-1 cells were stimulated with increasing amounts of either **A)** recombinant IFNγ, **B)** recombinant TNFα, or **C)** recombinant IL-2 for 1 hr prior to stimulation with cGAMP (0.5 µg). The resulting production of type I IFN in the supernatant was determined after 6 hrs of stimulation. Data in A) and B) show mean +SD and mean of duplicates from three independent experiments. Data in C) show mean of two duplicates from one experiment.

**Supplementary figure 3. T-EVs are not toxic to cancer cells *in vitro*.** MC38 cells were treated with murine T-EVs ($1 \times 10^9$) for 1 hr prior to stimulation with cGAMP (0.05µg) and compared to cells treated with 0.1µM doxorubicin. After 48 hrs, the viability of cells was determined using the CellTiter-Glo luminescent assay. Data show mean +SEM of 3 replicate samples.

**Supplementary figure 4. T-EVs alone have no anti-tumoral function in MC38 adenocarcinoma.** MC38 tumor bearing mice were treated with different amounts of T-EVs administered intratumorally (IT) 2 times with 3 days interval as indicated with black arrows on the figurers. Treatment started on day 9 after tumor cell inoculation. Data show tumor volume of each individual mouse from figure 6C, up to termination of the experiment at day 19.

**Supplementary figure 5. The anti-tumoral function of cGAMP is dose-dependent.** MC38 tumor bearing mice were treated with different amounts of cGAMP, as indicated, administered intratumorally (IT) 3 times with 3 days interval indicated with black arrows on the figurers.

Treatment started on day 9 after tumor cell inoculation. Data show tumor volume of each individual mouse from figure 6D, up to termination of the experiment at day 23. N=6 pr. group.

**Supplementary figure 6. T-EVs enhance antitumor efficacy of low-dosage cGAMP.** MC38 tumor bearing mice were treated with either T-EVs alone ($1.5 \times 10^8$), cGAMP alone (1µg) or a combination. Mice were treated IT 3 times with 3 days interval as indicated with black arrows on the figurers, starting on day 9 after tumor cell inoculation. Data show mean +/-SEM of tumor volume in mice treated with either Vehicle (EV-free control media, n=9), T-EVs alone (n=9), cGAMP alone (n=9), or a combination of T-EVs and cGAMP (n=8) up to day 15 after tumor cell inoculation. Data show tumor volume of each individual mouse from figure 6E, up to termination of the experiment at day 23.

**Supplementary figure 7. T-EVs combined with high-dosage cGAMP induce complete tumor regression.** MC38 tumor bearing mice were treated with either T-EVs alone ($1.5 \times 10^8$), cGAMP alone (10µg) or a combination. Mice were treated IT 2 times with 2 days interval as indicated with black arrows on the figurers, starting on day 10 after tumor cell inoculation. Data show tumor volume of each individual mouse from figure 6G. CR = complete responder.

### Supplementary figure 1

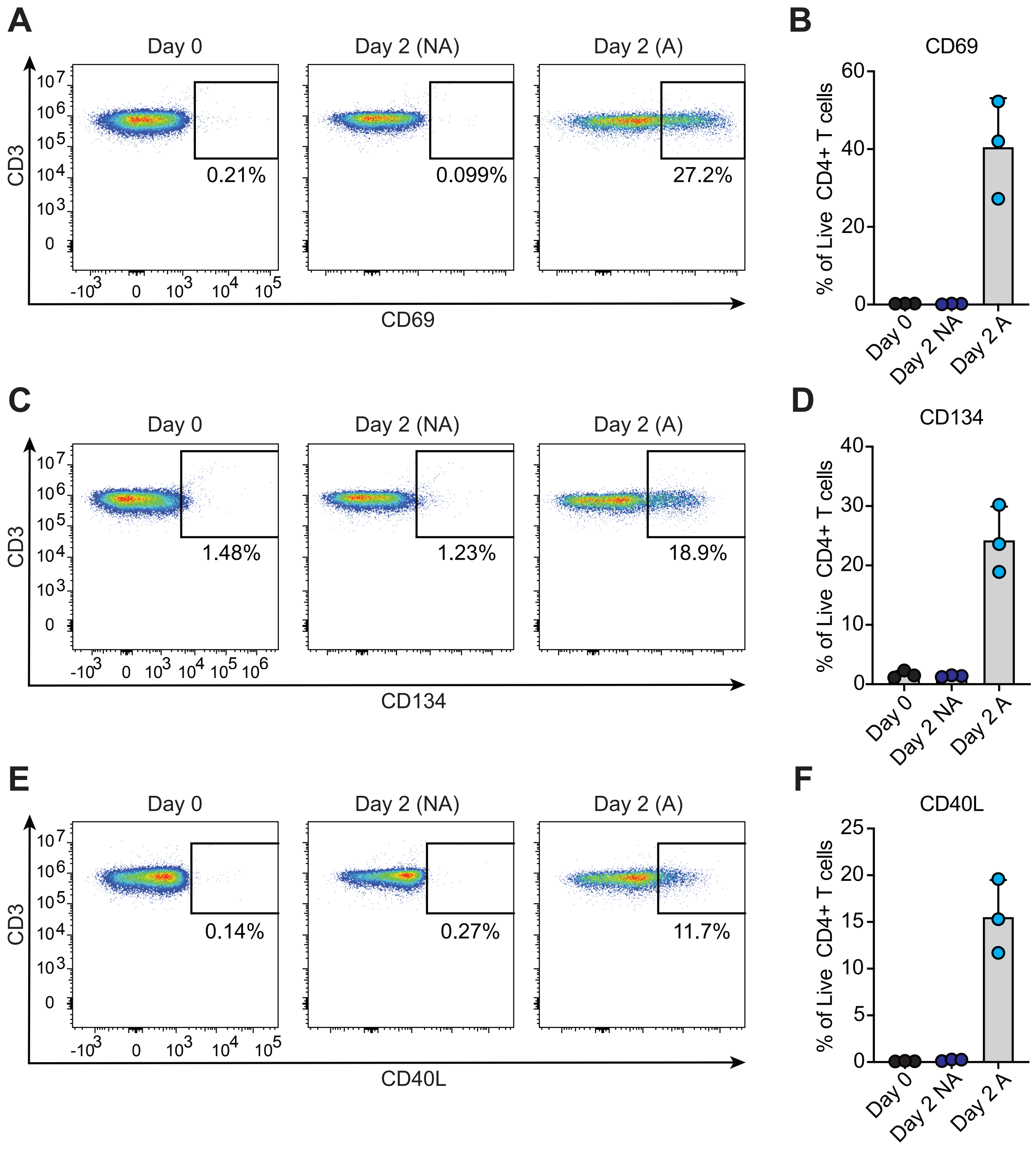

### Supplementary figure 2

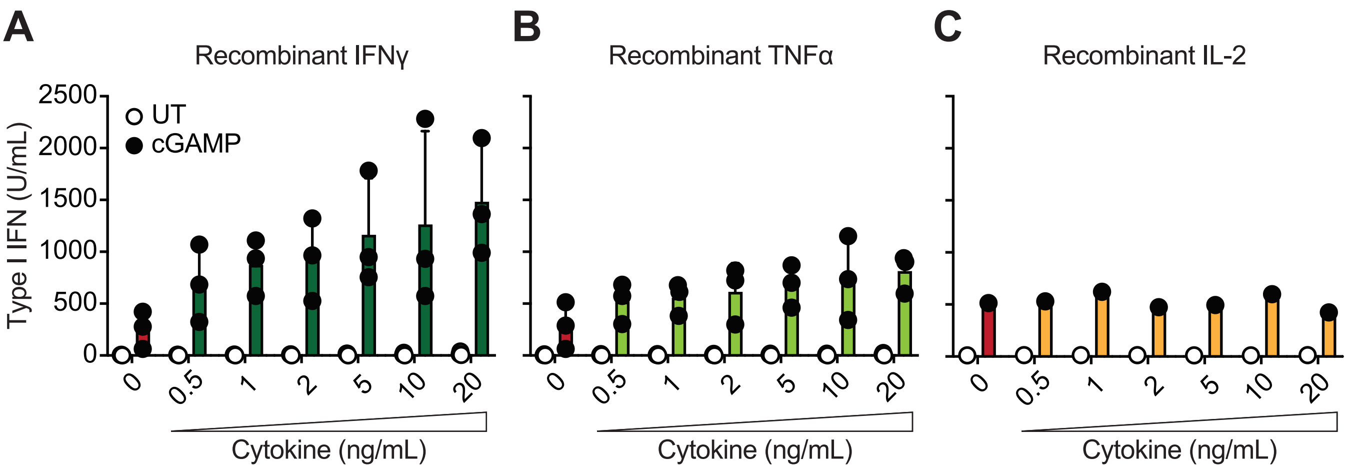

### Supplementary figure 3

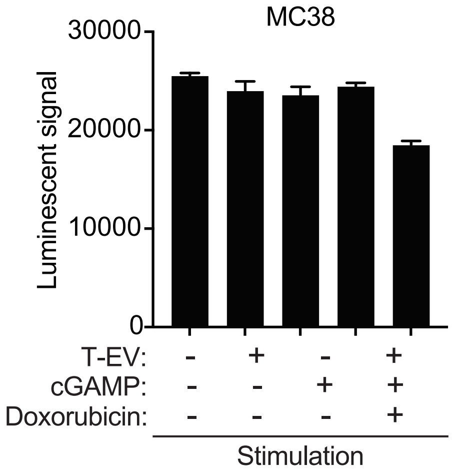

### Supplementary figure 4

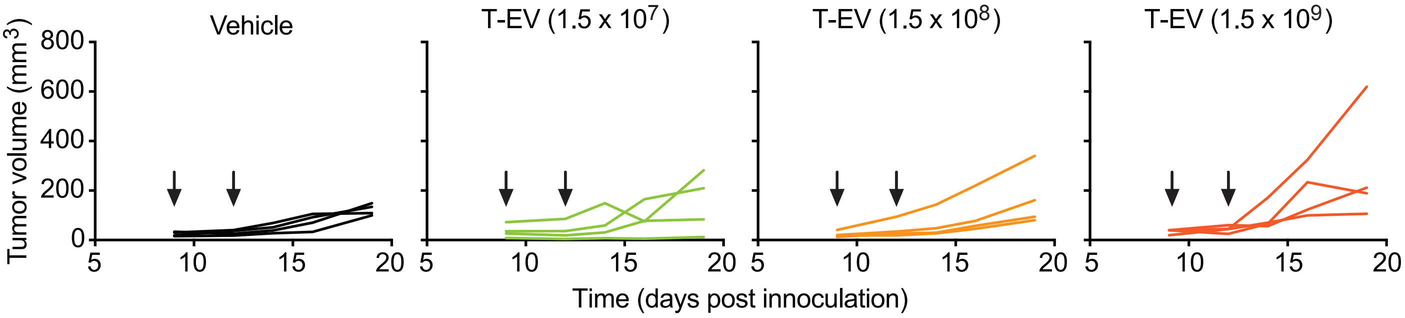

### Supplementary figure 5

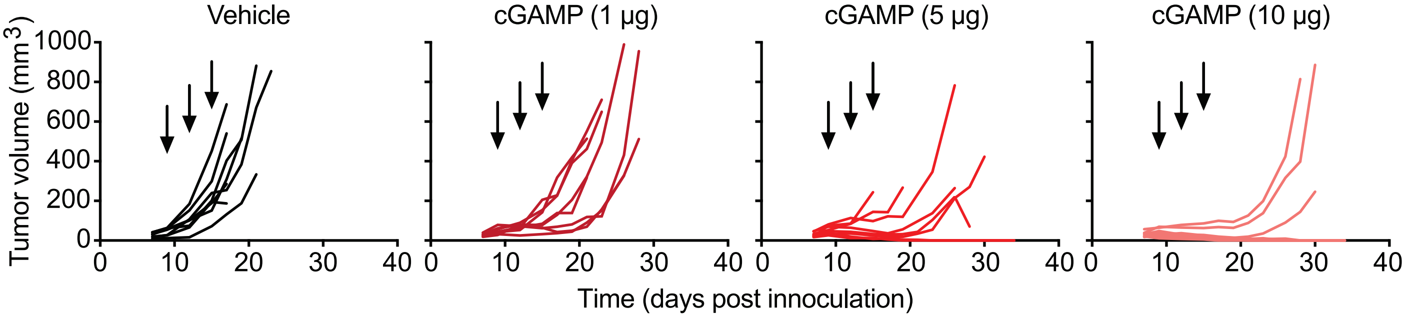

### Supplementary figure 6

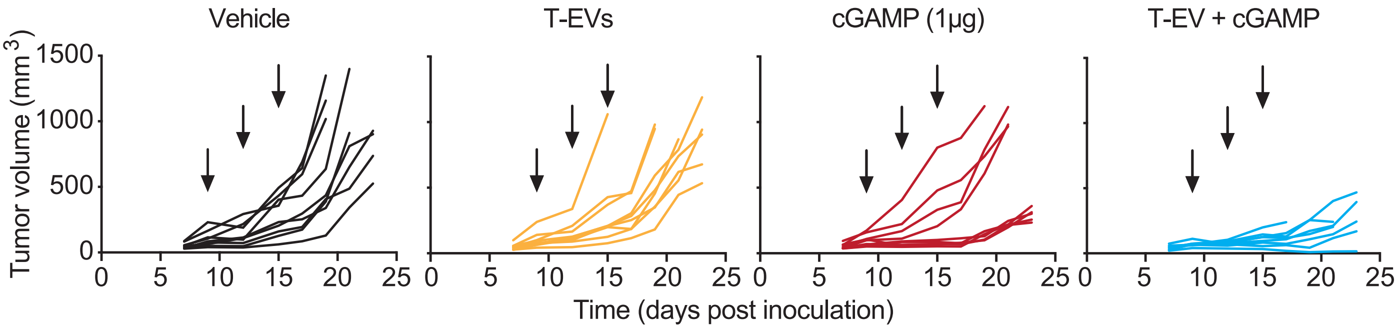

### Supplementary figure 7

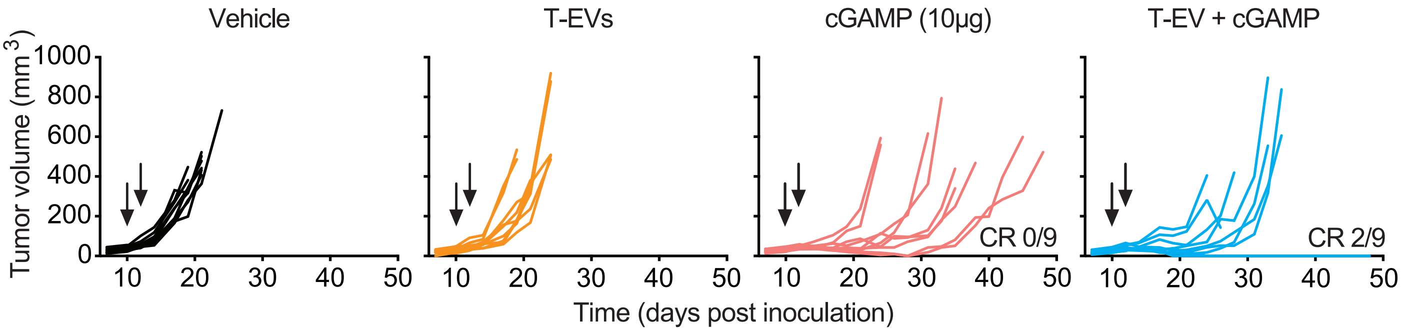
